# Direct estimation of genotype fitness from time series

**DOI:** 10.64898/2026.07.18.739367

**Authors:** Vaibhav Mohanty, Eugene. I. Shakhnovich

**Affiliations:** Department of Chemistry and Chemical Biology, Harvard University, Cambridge, MA 02138; Harvard/MIT MD-PhD Program, Harvard Medical School, Boston, MA 02115 and Massachusetts Institute of Technology, Cambridge, MA 02139; Program in Health Sciences and Technology, Harvard Medical School, Boston, MA 02115 and Massachusetts Institute of Technology, Cambridge, MA 02139

**Keywords:** fitness inference, fitness estimation, fitness landscape, time series, genotype correlations

## Abstract

Heterogeneous adapting populations, whether in laboratory evolution experiments or global-scale pandemics, experience complex evolutionary dynamics due to the interplay of selection, mutation, and stochasticity. Inference of individual genotypes’ fitnesses therefore becomes difficult, especially when many lineages are competing and data are noisy. Existing fitness inference methods tend to rely on assumptions on the fitness landscape’s maximum order of epistasis, or they require complicated iterative optimization algorithms to converge on fitness estimates. Here, we show that fitness landscapes can be computed from time series data, without any restrictions on epistatic order or iterative optimization, using a simple, closed-form mathematical expression that is easily implemented with standard matrix operations used commonly in linear algebra. We demonstrate successful fitness inference from noisy in silico evolutionary dynamics from four different noisy microscopic processes, including Wright-Fisher, Moran, ProSeD (serial dilution), and barcoded passage simulations. Then, we illustrate the broad applicability of the equation to five experimental time series datasets, including barcoded yeast evolution experiments, murine norovirus-1 serial passage experiments, and SARS-CoV-2 global genomic prevalence data. Our formula successfully infers fitnesses for even for rare genotypes several orders of magnitude less prevalent than top lineages, works with both laboratory evolution and epidemiological data, and can be implemented in most modern scientific programming languages.

## Introduction

Evolving populations climb toward the top of fitness landscapes semi-stochastically, exhibiting complex dynamics resulting from a combination of selection, mutation, demographic and environmental noise, and more (Wright 1932; Ewens 2004; de Visser and Krug 2014; Guo *et al*. 2019; Greenbury *et al*. 2022). Advances in high-throughput sequencing, including barcode sequencing, short- and long-read sequencing, and metagenomic sequencing, have enabled high-resolution tracking of lineage abundances over time, for eukaryotic, bacterial, and viral populations in the laboratory (Giallonardo *et al*. 2014; Blundell and Levy 2014; Levy *et al*. 2015; Good *et al*. 2017; Rotem *et al*. 2018; Nguyen Ba *et al*. 2019; Jasinska *et al*. 2020; Roodgar *et al*. 2021; Amato *et al*. 2022). These techniques, in addition to global efforts to collate pathogen genomic sequencing data, have led to the curation of large datasets of sequences such as GISAID (Elbe and Buckland-Merrett 2017; Shu and McCauley 2017; Wallau *et al*. 2023) with both geographic and temporal resolution. These experimental and epidemiological time series provide insight into the evolutionary properties of different genotype variants, such as mutational effects on strain or species fitness.

Fitness, classically defined as growth rate (Crow and Kimura 1970), is easily inferred from time series data when evolutionary competition is simple, like in the case of two large populations of mutants competing head-to-head in a serial passage simulation (Figure 1A). In this case, the species or strain with higher fitness (red) overtakes the other (blue) over time with a sigmoidal frequency trajectory. The fitness difference between two strains for such uncomplicated time series is easily predicted by decades-old results from classical population genetics (Watterson 1982). But realistic evolutionary dynamics, both in laboratory evolution experiments and for globally prevalent pathogens, are typically far more complicated (Figure 1B). Often, many strains compete at the same time (clonal interference), random mutations are constantly taking place in all lineages, and noise enters the time series from many sources, including genetic bottlenecks, environmental factors, sampling of sequences for measurement, and measurement errors (Gerrish and Lenski 1998; Desai and Fisher 2007; Fogle *et al*. 2008; Blundell and Levy 2014; Levy *et al*. 2015; Mohanty *et al*. 2026).

**Figure 1.**
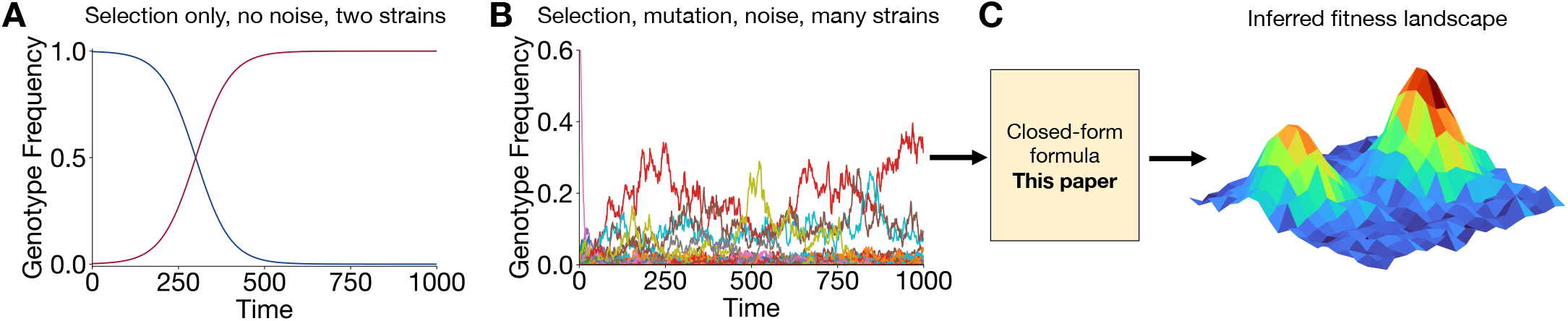
Complexity of evolutionary trajectories and inference of fitnesses from genotype time series. (**A**) Simple competition between two competing lineages in the infinite population limit follows logistic functions from which fitness difference can be easily inferred. (**B**) In general, selection, mutation, and genetic drift (demographic noise) create complex evolutionary trajectories with genotype frequencies spanning potentially several orders of magnitude. (**C**) Here, we introduce a simple, closed-form formula to infer fitnesses for observed genotypes even for complex evolutionary trajectories.

Several methods, often specific to the type of sequencing performed, have emerged to infer lineage fitnesses from time series data (Bollback *et al*. 2008; Illingworth and Mustonen 2011; Malaspinas *et al*. 2012; Mathieson and McVean 2013; Shekhar *et al*. 2013; Feder *et al*. 2014; Lacerda and Seoighe 2014; Steinrücken *et al*. 2014; Foll *et al*. 2015; Topa *et al*. 2015; Terhorst *et al*. 2015; Illingworth 2015; Schraiber *et al*. 2016; Ferrer-Admetlla *et al*. 2016; Gompert 2016; Aita *et al*. 2016; Barton *et al*. 2016; Iranmehr *et al*. 2017; Taus *et al*. 2017; Li *et al*. 2018; Zinger *et al*. 2019; Liu *et al*. 2019; Nguyen Ba *et al*. 2019; Paris *et al*. 2019; He *et al*. 2020; Zeng and Aurell 2020; Sarkisov *et al*. 2020; Salehi *et al*. 2021; Sohail *et al*. 2021, 2022; Obermeyer *et al*. 2022; Li *et al*. 2023; Li and Barton 2024; Sundar *et al*. 2025; Zeng *et al*. 2025). Some methods focus on estimating fitness from single allele frequencies, while others aim to build an epistatic picture of the fitness landscape using long-read sequencing data. Every method has its own shortcomings. For example, many methods which predict epistatic fitness landscapes from full genotype frequencies rely on the assumption that the maximum order of epistatic interactions is capped, effectively projecting higher order epistatic interactions onto a fitness model which includes only up to pairwise interactions between loci (e.g. Sohail *et al*. (2022); Obermeyer *et al*. (2022)). Other methods make no prior assumptions about the structure of the fitness landscape but limit the number of time steps in the series (Sundar *et al*. 2025) or base their inference pipeline on details specific to their experimental setup, leading to complex, probabilistic models that require iterative optimization to converge on fitness estimates (e.g. Nguyen Ba *et al*. (2019); Li *et al*. (2018, 2023)). These methods also do not handle cases when mutation is not weak compared to selection.

Here, we report that fitnesses can be inferred from genotype frequency time series data using only a simple, closed-form mathematical expression, which can be implemented using common matrix operations found in standard scientific programming languages, or even simply using spreadsheet software. Our equation follows directly from classical population genetics and works on a variety of time series datasets, even when mutations are not weak compared to selection. The computational pipeline (Figure 1C) needed to infer fitnesses is simply to transform the frequency time series into a particular matrix form, average over all timepoints, and then compute the pseudoinverse of the matrix, typically pre-implemented in numerical linear algebra packages using singular value decomposition (SVD), which is widely used for data compression and signal processing (Van Der Veen *et al*. 1993). Modern programming languages used widely in science such as Python, R, MATLAB, and Julia contain such pre-implemented methods, or the user can even utilize our pipeline without any programming, in Microsoft Excel.

To test the accuracy and validity of the method, we first apply the equation to four types of evolutionary dynamic simulations, including the standard Wright-Fisher process (Ewens 2004; Greenbury *et al*. 2022; Good 2024), standard Moran process Moran (1962); Ewens (2004); Good (2024), microbial serial dilution simulation (ProSeD algorithm) (Mohanty *et al*. 2026), and barcoded cell growth simulation (Fit-Seq2.0 Growth Simulation) Li *et al*. (2023). Then, to showcase the broad applicability of the equation to different experimental and epidemiological data sets, we apply the equation to infer fitnesses from five empirical genotype time series datasets, including yeast barcode evolution experiments in two different growth media Nguyen Ba *et al*. (2019), murine norovirus 1 (MNV-1) bulk serial passage experiments with and without a neutralizing antibody Rotem *et al*. (2018), and global genomic sequencing data for SARS-CoV-2 Elbe and Buckland-Merrett (2017); Shu and McCauley (2017); Wallau *et al*. (2023). For the simulated datasets, we find excellent agreement between our inferred fitnesses and ground-truth fitnesses, even for rare genotypes whose average prevalence was orders of magnitude lower than the most frequently observed lineages. For the experimental datasets, we find that our inferred fitnesses are strongly correlated with those inferred from other methods, including ones which require iterative optimization.

In this work, we show that classical population genetics admits a mathematically simple solution to the fitness inference problem that is valid across a wide array of simulated and experimental evolution datasets. The formula provides comparable performance to other fitness inference methods, as evidenced by similar performance on experimental and simulated datasets. While other existing methods may offer specific benefits for their target use cases, we emphasize that simplicity of our reported formula, and its computational implementation, offer a convenience and ease of use that should provide utility to evolutionary biologists, microbiologists, and population health scientists alike.

## Results

### Simple, closed-form formula for fitness inference from genotype time series

#### Analytical formula

Consider a population of asexual, haploid individuals evolving in a setting where population size *N* is fixed, and suppose that there are *G* genotypes which may be observed. The multi-genotype generalization of the well-known Kimura equation (Crow and Kimura 1970) captures how the frequency *f*_*g*_ (*t*) of a genotype *g* is impacted over time by the interplay of natural selection, mutation, and genetic drift (stochastic fluctuations due to discrete inter-generation sampling):

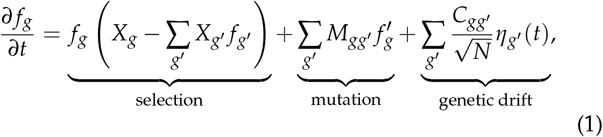

where *X*_*g*_ is the fitness of genotype *g, M* is a mutation rate matrix with off-diagonal elements *M*_*gg*_′ = *µ*_*g*→*g*_′ for *g*≠ *g*′ and diagonal elements *M*_*gg*_ = − ∑_*g*_′_≠*g*_ *µ*_*g*_′ _→*g*_, *δ*_*g,g*_′ is the Kronecker delta, and 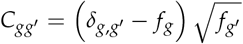 is the element of the matrix which sets the covariance structure of the white noise terms *η*_*g*_′ (*t*). Working in the limit of infinite population (*N* → ∞), we then time-average both sides of the equation and rearrange to solve for the vector of fitness estimates **X** = (*X*_1_, … , *X*_*G*_), as the central result of this work, which is a simple and closed-form formula that infers relative Malthusian fitness up to a linear shift:

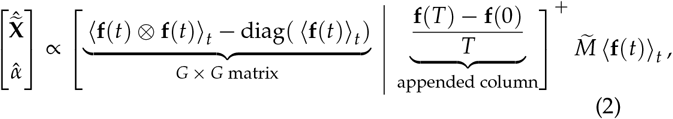

Above, **f**(*t*) = (*f*_1_(*t*), … , *f*_*G*_ (*t*)) is the frequency time series as a function of time t which runs from *t* = 0 to *t* = *T*, the hats *ŷ* on a value *y* indicates its inferred estimate, 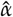 is a free positive constant that can be discarded, ⟨·⟩_*t*_ indicates time-averaging, diag(**y**) takes a vector **y** and places it on the diagonal of a matrix, **a** ⊗ **b** is the outer product of two vectors **a** and **b**, the superscript “+” indicates the matrix pseudoinverse operation, and 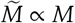 is proportional to the known or hypothesized mutation matrix. A complete derivation can be found in Supplementary Information Section S1.

#### Computational implementation

Practical implementation of the formula is straightforward. Storing the frequency time series **f**(*t*) as a *G* × (*T* + 1) matrix (since time steps run from 0 to *T*), we first compute the column vector (**f**(*T*) − **f**(0))/*T*, which is the average change in frequencies across the time interval. Then, we compute the time average ⟨**f**(*t*) ⟩_*t*_ by averaging over columns in the *G* × (*T* + 1) matrix. Next, we compute the outer product **f**(*t*) ⊗ **f**(*t*) for each time point *t*, resulting in a *G* × *G* × (*T* + 1) array, which is time-averaged across the time dimension to result in a *G* × *G* matrix. Placing ⟨**f**(*t*)⟩_*t*_ on the diagonal of a *G* × *G* matrix, we can easily construct ⟨**f**(*t*) ⊗ **f**(*t*) ⟩_*t*_ − diag(⟨**f**(*t*)⟩_*t*_)), and append (**f**(*T*) − **f**(0))/*T* to the right of the matrix, resulting in a *G* × (*G* + 1) matrix. The pseudoinverse of this matrix can be calculated using SVD; the smallest singular values can be filtered, if desired. Linear algebra packages in common programming languages frequently have pseudoinverse methods pre-implemented, such as NumPy’s (Harris *et al*. 2020) `numpy.linalg.pinv`.

The last step is to calculate the mutation rate matrix 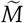. If the mutation rate matrix is exactly known, with respect to the correct units of time, then an alternate formula (Supplementary Information Section S2) can be used. If mutations are exactly zero, then eq. (2) cannot be directly used, an another alternate formula (Supplementary Information Section S3) can be used instead. The latter formula works well for evolution which takes place in the strong selection, weak mutation (SSWM) regime, and in fact it *provably* provides Pearson correlation of 1 between estimated and ground truth fitness for a series of selective sweeps, which we rigorously show in Supplementary Information Section S3.

In general, however, mutation rates are not known, but genotype sequences may be known; in these cases, we recommend hypothesizing 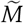 by estimating the probability of point mutations, up to a multiplicative constant; we will demonstrate how this can be successfully used for inferring SARS-CoV-2 fitness from global genomic sequencing data. Finally, some experimental evolution settings (such as barcoding experiments with many possible barcodes, or large genomes with long-read sequencing) have intractably large or complex genotype spaces. In these cases, the vast majority of possible genotypes go unobserved in the frequency time series **f**(*t*), meaning their average frequencies ⟨**f**(*t*)⟩ are zero (or close to it). A direct consequence of this is that, for an observed genotype *g*, the vector element 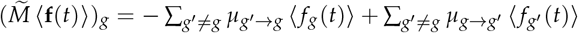 will have far stronger dependency on the contribution of *outgoing* mutations − ∑_*g*_′_≠*g*_ *µ*_*g*_′ _→*g*_ ⟨*f*_*g*_ (*t*)⟩ than *incoming* mutations ∑_*g*_′_≠*g*_ *µ*_*g*→*g*_′ ⟨ *f*_*g*_′ (*t*)⟩, since large number of ⟨*f*_*g*_′ (*t*)⟩ values will be zero for unobserved genotpyes *g*′. This permits replacing the scaled mutation matrix 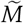 with a negative identity matrix 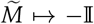. A complete derivation of this simplifying approximation is shown and justified in Supplementary Information Section S4. Additionally, we will show that this approximation is justified *a posteriori* by its performance on barcoding simulations, laboratory barcoded yeast evolution experiments, laboratory MNV-1 serial passage experiments later in this work.

Fitness estimates, up to proportionality, can then be estimated by substituting the constructed matrices and vectors into eq. (2). These steps can be performed in any common scientific programming language or even in Microsoft Excel. Accuracy of the fitness estimates will depend on presence of fitness data: naturally, vanishingly rare genotypes will have low signal-to-noise ratio and thus inaccurate fitness estimates. We thus recommend using all filtering fitness estimates by retaining genotypes only above a threshold average frequency, which we will refer to as the “average (genotype) frequency cutoff ⟨*f* (*t*) ⟩ _*t*_” going forward.

### Validation: simulated evolutionary dynamics

To assess the accuracy of the fitness estimation formula, we simulated evolutionary dynamics on fitness landscapes where ground-truth fitnesses are known. Four different microscopic processes were simulated: (1) the standard Wright-Fisher process where each generation is obtained by fitness-weighted random sampling of the previous generation (Ewens 2004; Greenbury *et al*. 2022; Good 2024), (2) a variant of the Moran process in which two individuals compete to replace each other at each timestep as described by Good (2024), (3) ProSeD, a microbial serial dilution simulation with overlapping generations recently introduced by Mohanty *et al*. (2026), and (4) the barcoded cell Growth Simulation introduced in parallel with Fit-Seq2.0 by Li *et al*. (2023). The numerical details for these simulations are provided in the Materials and Methods. Example genotype frequency trajectories for each of these simulation types, shown in Figure 2A, demonstrate complex and stochastic trajectories for many genotypes whose average frequencies can span several orders of magnitude—e.g. ≈ 5 orders of magnitude for 1000 unique genotypes (Figure S1A).

**Figure 2.**
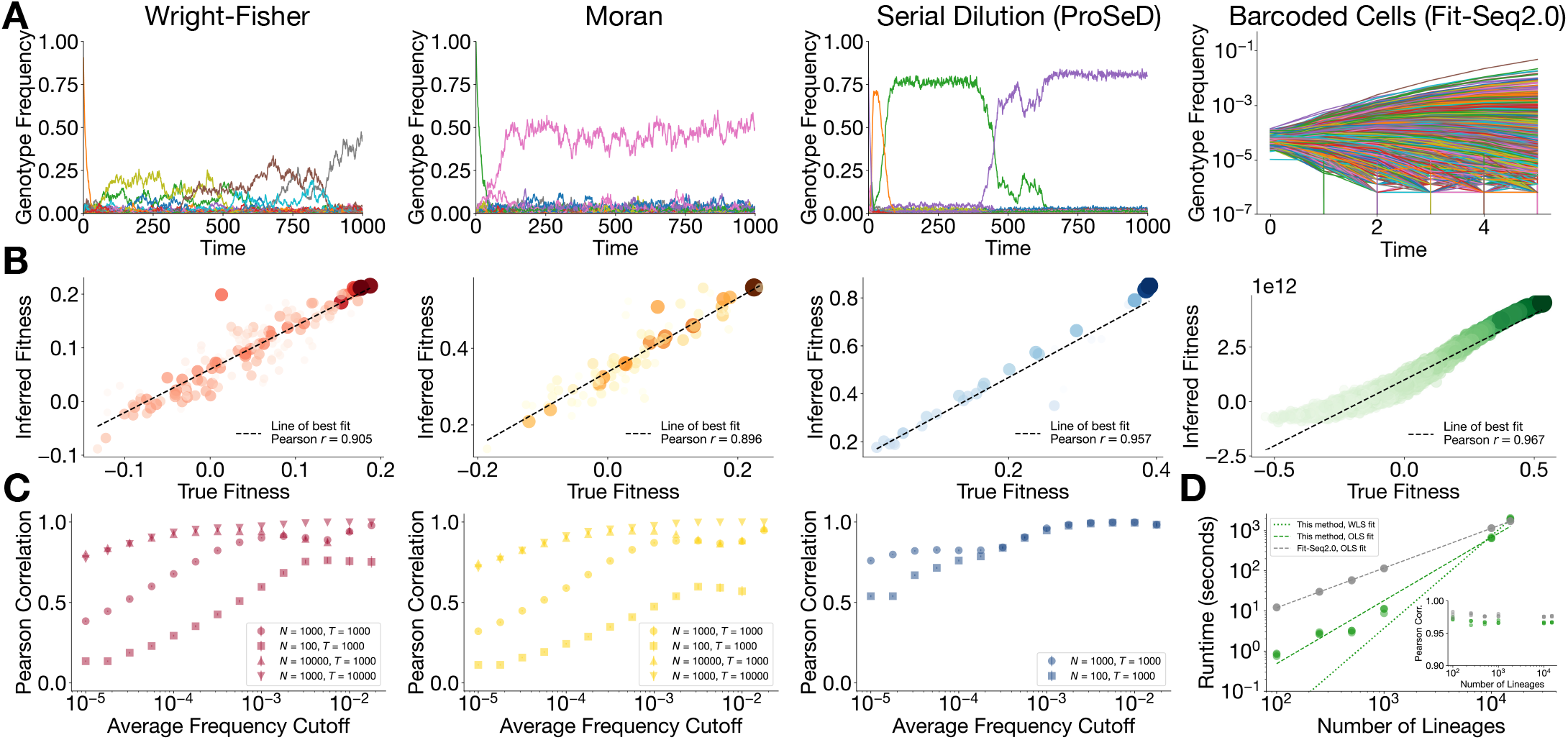
Simulated evolutionary dynamics and accurate fitness inference using eq. (2). (**A**) Example genotype frequency time series for Wright-Fisher, Moran, ProSeD, and Fit-Seq2.0 Growth Simulations. Trajectories can be noisy and complex, with orders of magnitude variation in genotype average frequency. (**B**) Comparison between ground truth fitnesses and fitnesses inferred using eq. (2) on example trajectories for each simulation type, evaluating on genotypes whose average frequency was ⟨*f* (*t*) ⟩_*t*_ ≥ 10^−3^. Equation (2) demonstrates faithful reconstruction of the observed fitnesses even for rare genotypes. (**C**) Pearson correlation between ground-truth fitnesses and fitnesses inferred using eq. (2), as a function of average frequency cutoff. Pearson correlation for multiple population sizes and/or simulation lengths are shown using different symbols; error bars show standard errors from averaging over 100 trials. (**D**) Runtime of the Python implementation of eq. (2) and of Fit-Seq2.0 versus number of lineages, demonstrating power-law scaling for both. Inset: Pearson correlation between ground-truth and inferred fitnesses for both inference methods, plotted versus number of lineages.

We ran Wright-Fisher, Moran, and ProSeD simulations on random epistatic fitness landscapes (Materials and Methods) for multiple population sizes (*N* = 10^2^ to 10^4^) and/or multiple time ranges (*T* = 10^3^ or 10^4^), and the Fit-Seq2.0 Growth Simulation was run for population size *N* = 10^4^ for 5 broth transfers. Time series data were recorded from these simulations and used to infer fitnesses of the top frequent genotypes using eq. (2); these were subsequently compared to the known ground-truth fitnesses. For the Wright-Fisher, Moran, and ProSeD simulations, the mutation rate matrices were exactly known, so we utilized them directly in eq. (2), and for the Fit-Seq2.0 Growth Simulation data we used the negative identity matrix.

Following the protocol described in the previous section, we use all genotypes in the timeseries to infer fitness, but we evaluate performance of eq. (2) for subsets of genotypes filtered by different average frequency cutoffs. Results of fitness inference for an example trajectory from each of the four simulation types are shown in Figure 2B; the Wright-Fisher, Moran, and ProSeD simulations use an average frequency cutoff of ⟨*f* (*t*)⟩ _*t*_ ≥ 10^−3^, while all genotypes are retained for the Fit-Seq2.0 Growth Simulation. Each of these demonstrates high accuracy in reconstructing the fitness landscape, with Pearson correlations ranging from near ≈ 0.90 to ≥ 0.96, depending on the simulation. In Figure S1B, an average frequency cutoff of ⟨*f* (*t*)⟩ _*t*_ ≥ 10^−4^ is used to infer fitness from the same Wright-Fisher, Moran, and ProSeD simulation trajectory, showing, as expected, that more genotypes’ fitneses are inferred with a lower frequency cutoff, but the overall Pearson correlation between inferred and true fitnesses decreases compared to the ⟨ *f* (*t*)⟩_*t*_ ≥ 10^−3^ threshold.

Figure 2C shows the relationship between Pearson correlation (measuring inference performance) and average frequency cutoff for the Wright-Fisher, Moran, and ProSeD simulations, averaged over 100 simulation trials. These 100 trials are repeated for different population sizes and/or simulation time lengths. All three simulation types show common trends: Pearson correlation between inferred and true fitness tends to *increase* with a higher average frequency cutoff (since fewer, more frequent genotypes’ fitnesses are being inferred), with increasing population size (since the effect of noise from genetic drift decreases for larger populations), and with increasing simulation time (which provides more data points for averaging). Figure S1C yields similar conclusions, but with Pearson correlation plotted against the *number* of genotypes inferred, which is also set by the average frequency cutoff. Naturally, fewer genotypes are inferred as the average frequency cutoff is raised, and inference becomes less noisy and more accurate. We also ran Wright-Fisher simulations for a wide range of mutation probabilities spanning several orders of magnitude. We found that Pearson correlation for fitness estimation did not change much at any given average frequency cutoff for sufficiently low mutation probabilities; very high mutation probabilities led to rapid spreading of the population across the entire genotype space, but these cases are unlikely experimental scenarios (Figure S2A). Changing mutation probabilities did, however, change how many fitnesses were inferrable at a given accuracy level, as measured by Pearson correlation; this is because decreased mutation probabilities also decreased the number of observed fitnesses at a given average frequency cutoff (Figure S2B).

Ultimately, population size, simulation time, and mutation rate had expected and intuitive impacts on the accuracy of fitness estimation: larger population size and longer simulation times improved Pearson correlation at a given frequency cutoff and allowed for more fitnesses to be inferred simulatneously, while increasing mutation rate did not change Pearson correlation at a given frequency cutoff but did allow for more fitnesses to be inferred at high accuracy due to increased exploration of the genotype space. In general, the simulated evolutionary dynamics demonstrate that known ground-truth fitnesses can be inferred up to linear transformation with high accuracy using eq. (2), with the accuracy correlating positively with larger population size, longer simulation times, and higher average frequency cutoffs, which also implies fewer genotypes inferred. A principal conclusion to highlight is that, despite highly noisy trajectories and genotypes’ average frequencies spanning several orders of magnitude, eq. (2) shows that an infinite-population assumption still works well for fitness inference; we attribute this recoverability of the fitness information to the genotype-genotype correlations which are encoded into the *G* × *G* matrix in eq. (2).

#### Recommended number of lineages for fitness estimation based on runtime and comparison to extant method

We compared the runtime of our Python implementation of eq. (2), which relies on NumPy’s SVD-based pseudoinverse function, to Fit-Seq2.0’s inference pipeline (Li *et al*. 2023) on simulated evolutionary trajectories from the Fit-Seq2.0 Growth Simulation. Setting the number of lineages *N*_*g*_ to values between 10^2^ and 1.5 × 10^4^, we measured the runtime *τ* of both fitness estimation algorithms on the same machine (MacBook Pro, Apple M1 Pro chip, 32 GB memory) over three trials. Both eq. (2) and Fit-Seq2.0 demonstrate power-law scaling with the number of lineages 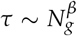 (Figure 2D): Fit-Seq2.0 has a power-law exponent *β* ≈ 0.989 (close to linear scaling), while eq. (2) has power-law exponent *β* ≈ 1.57 (as measured by ordinary least squares fitting) or *β* ≈ 2.30 (as measured by weighted least squares fitting with weights proportional to number of lineages). The Python implementation of eq. (2) uses SVD, which has a known complexity of *O*(*n*^3^) for *n* × *n* matrices (Golub and Van Loan 2013), so in principle our method should have worst-case performance *β* ≈ 3.

Empirically, we see though that until the number of lineages reaches close to 15,000, the Python implementation of eq. (2) runs faster than Fit-Seq2.0 (Figure 2D); for more lineages, Fit-Seq2.0’s linear scaling gives it the advantage. Nonetheless, for all cases we studied, eq. (2) and Fit-Seq2.0 achieve comparable Pearson correlation (≥ 0.95) on fitness inference for all lineages, with Fit-Seq2.0 carrying a slight accuracy advantage while eq. (2) carries a speed advantage for fewer than approximately 15,000 lineages. Overall, we thus recommend using eq. (2) as a rapid fitness estimation method if the number of lineages for which fitness is inferred is fewer than roughly 15,000 since it carries comparable accuracy, performs faster, and—being a single equation—is considerably simpler to implement.

### Validation: experimental and epidemiological time series

Now, we demonstrate the applicability of eq. (2) to a variety of evolutionary datasets: barcoded yeast experiments, MNV-1 serial dilution viral evolution experiments with long-read sequencing, and SARS-CoV-2 global genomic sequence prevalence data. Nguyen Ba *et al*. (2019) performed barcoded yeast evolution experiments, collecting time series for budding yeast lineages in two different media: yeast extract peptone dextronse (YPD) and yeast extract peptone acetate (YPA). They obtained 110 sequencing time points (once every 24 hours, roughly 10 generations) for the YPD medium evolution and 99 sequencing time points for the YPA medium evolution. We (Rotem *et al*. 2018) previously performed MNV-1 serial dilution experiments with (“Ab+”) and without (“Ab-”) monoclonal antibody A6.2, sequencing every 24 hours, and obtaining five sequencing time points for each antibody conditions. Finally, we (Wang *et al*. 2024) previously collected SARS-CoV-2 genomic prevalence data from the GISAID database for a subset of strains in which the spike protein receptor binding domain (RBD) was mutated between wildtype and Omicron SARS-CoV-2 strains, representing over 3 years of prevalence data; here, we binned the prevalence data into 300 equally spaced time points, obtaining genotype frequency time series over this time domain. All genotype frequency time series are shown in Figure 3A; genotype average frequencies span roughly 4-5 orders of magnitude (Figure S3A).

**Figure 3.**
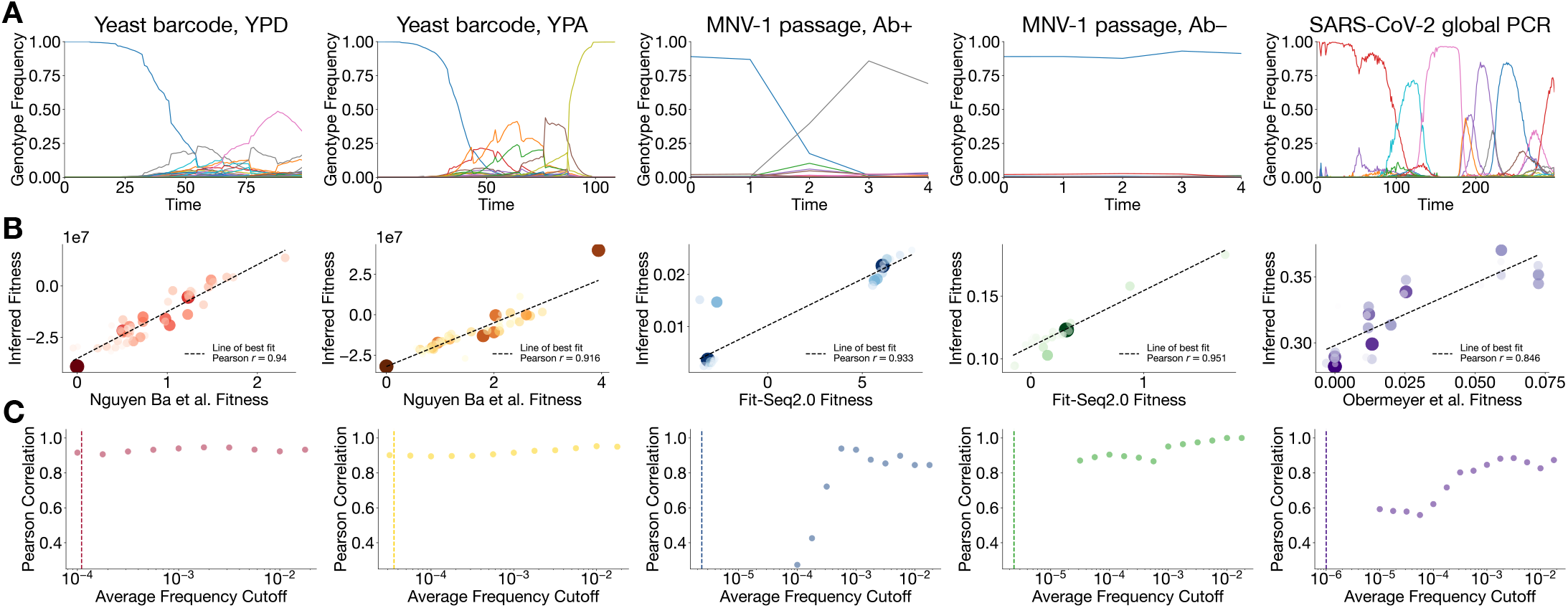
Empirical evolutionary dynamics from *in vitro* experiments and global genomic prevalence data and fitness inference using eq. (2). (**A**) Genotype frequency time series from barcoded yeast evolution experiments (YPD, YPA), MNV-1 evolution experiments (Ab+ and Ab−), and SARS-CoV-2 genomic prevalence. (**B**) Comparison between fitnesses inferred using eq. (2) and fitnesses inferred by other methods in the literature; we included genotypes whose average frequency was ⟨*f* (*t*)⟩ _*t*_ ≥ 10^−3^. Equation (2) demonstrates reconstruction of the observed fitnesses comparable to those found by other methods. (**C**) Pearson correlation between fitnesses inferred using eq. (2) and literature-reported fitnesses, as a function of average frequency cutoff.

Unlike the simulated evolutionary trajectories, ground truth fitnesses were not known for these five datasets. So, we compared our fitness estimates to fitnesses inferred via other methods, whether by the original authors (if available) or by us. Nguyen Ba *et al*. (2019) developed their own iterative optimization method for fitness inference and provided reported values for the two barcoded yeast evolution experiments. For our previous MNV-1 evolution experiments (Rotem *et al*. 2018), we estimated fitnesses of individual variants using Fit-Seq2.0 (Li *et al*. 2023) in a recent work (Mohanty and Shakhnovich 2026). SARS-CoV-2 fitnesses were obtained from Obermeyer *et al*. (2022), which we had previously compiled (Wang *et al*. 2024). The SARS-CoV-2 fitnesses from Obermeyer *et al*. (2022) assume a completely additive fitness landscape, while the yeast fitnesses from Nguyen Ba *et al*. (2019) and the MNV-1 fitnesses from (Mohanty and Shakhnovich 2026) made no restrictive assumptions on the order of epistasis.

Here, we estimated fitness for these five datasets (YPD yeast, YPA yeast, MNV-1 Ab+, MNV-1 Ab− , SARS-CoV-2 genomic prevalence) using eq. (2). For yeast experiments, the barcode frequencies had been merged into lineages by the authors, so the lineage counts were simply normalized to sum to one at each time point to obtain lineage frequencies; for MNV-1, raw read counts were similarly normalized to one at each time point; and for SARS-CoV-2, genomic sequence hits within each time point were summed to one at each time point. For the yeast and MNV-1 time series, we used the approximation that the mutation matrix could be replaced by the negative identity matrix; as mentioned previously in the “Computational implementation” subsection and justified mathematically in Supplementary Information Section S4, this assumption should be used when the mutation matrix is not known or difficult to calculate, and the genotype space is very large—both of which apply to barcoded yeast evolution and MNV-1 serial evolution with long-read sequencing. For the SARS-CoV-2 timeseries, we constructed an approximated mutation matrix by estimating the average probability of amino acid transitions by assuming that the underlying trinucleotide codons had uniform point mutation probabilities.

Using an average frequency cutoff of ⟨*f* (*t*)⟩ _*t*_ ≥ 10^−3^, we find excellent Pearson correlation between fitnesses inferred with eq. (2) and the other extant inference methods, as seen in Figure 3B. As a supplementary reference, we also show a comparison between our inferred fitnesses and other methods for a lower average frequency cutoff ⟨*f* (*t*)⟩_*t*_ ≥ 10^−4.5^ (Figure S3B), which, as expected, shows a decrease in Pearson correlations. The Pearson correlation is shown as a function of the average frequency cutoff for all five datasets in Figure 3C. These panels show that the barcoded yeast experiments’ Pearson correlations are largely robust to average frequency cutoff, even when all data points are included. MNV-1 Pearson correlations decrease precipitously for average frequency cutoffs lower than 10^−3.25^ for Ab+ and 10^−4.5^ for Ab −; this is likely due to many new data points being included in the evaluation which had highly uncertain Fit-Seq2.0 fitnesses in the first place (see large error bars in Figure S3B). Unlike the yeast and MNV-1 cases, the SARS-CoV-2 fitness estimation Pearson correlation decreases with frequency cutoff gradually until plateauing near 0.6 between average frequency cutoffs between 10^−4^ to 10^−5^. Similar plots showing Pearson correlation and the number of genotypes whose fitnesses are inferred are provided in Figure S3C.

We emphasize that for all of these datasets, the true ground-truth fitness is not known. Therefore, too much emphasis should not be placed on the exact Pearson correlations between our inferred fitnesses and those from other methods, since every method carries its own simplifying assumptions, such (Obermeyer *et al*. 2022) assuming that the SARS-CoV-2 fitness landscape can be taken to be non-epistatic, where mutational effects additively contribute to fitness. Rather, we simply highlight that generally high correlations between our method and others demonstrates that salient fitness information can be extracted from the time series data conveniently, without iterative optimziation procedures like those employed by Nguyen Ba *et al*. (2019) or Fit-Seq2.0 (Li *et al*. 2023), and without simplifying assumptions on the degree of epistasis like in Obermeyer *et al*. (2022). The previous section with simulated evolutionary dynamics data already showed that the method is accurate when the dynamics are derived directly from known ground-truth fitnesses.

Moreover, the successful application of eq. (2) to infer fitnesses for five experimental time series datasets representing three unique types of lineage data (barcoded yeast evolution, viral evolution with long-read sequencing, and genomic prevalence data) demonstrates that the method we present here is flexible toward the data source. The data can be *in vitro* experimental data; the lineages may evolve due to acquired barcodes or random point mutations in *in vitro* settings; and even pathogen genomic prevalence data can simply be aggregated, binned in time, and normalized into genotype frequencies. Equation (2) therefore is comparable to other methods, does not restrict the type of data used to construct the frequency time series, and also enjoys ease of implementation.

## Discussion

In this work, we have introduced a simple analytical formula, eq. (2), following directly from the infinite population limit of the multilocus Kimura equation which permits the inference of fitness estimation from multi-genotype frequency time series data without restrictions on the degree of epistasis. In experimental settings, we showed concurrence between our method and extant methods, for three different types of genotype data including barcoded yeast evolution, long-read sequencing data from viral evolution, and global pathogen genomic sequence prevalence. To demonstrate accuracy with respect to known ground-truth epistatic fitness landscapes, we tested eq. (2) on simulated evolutionary dynamics from four different types of discrete-time, agent-based evolution algorithms, achieving high accuracy on fitness estimates for genotypes ranging across four to five orders of magnitude in average frequency. The key lies in the pseudoinversion of a matrix which captures genotype-genotype correlations over time; even rare genotypes’ fitnesses can be inferred because they interact with more frequent genotypes potentially over many time steps.

Implementation of eq. (2) introduced here, like any other method for fitness estimation, possess its own limitations. One is the asymptotic runtime scaling of eq. (2); because pseudoinversion of the matrix in eq. (2) is often implemented using SVD, the runtime scales as a supralinear power law, and the Python implementatino of eq. (2) becomes slower than Fit-Seq2.0 for fitness inference for more than ≳ 15,000 lineages. Another limitation is that eq. (2) does not provide error estimates on the fitnesess. Therefore, we have suggested filtering fitness estimates by retaining fitness estimates for genotypes whose time-averaged frequency is greater than a user-chosen cutoff. This cutoff is somewhat arbitrary, as different cutoffs lead to different accuracies depending on the shape of the fitness landscape, the source of the data, etc. However, we argue that this choice of frequency cutoff is no more arbitrary than the choice of threshold for a binary classifier, for example; in general, one can always sweep over the average frequency cutoff parameter to establish accuracy curves such as in Figure 2C and Figure 3C.

Equation (2) is primarily notable for its simplicity and ease of implementation. It can be implemented in any major scientific programming language which has access to standard linear algebra operations such as SVD; and the equivalent operations can even be performed in standard spreadsheet software. We believe it can serve as a useful alternative to current software which may have a higher barrier to usage due to software and dependency installation time. Experimentalists conducting laboratory evolution—such as the barcoded yeast evolution and viral evolution considered here—may benefit from the use of eq. (2) in estimating fitness from pooled competition experiments.

In the broader context in the field of population genetics, perhaps the most striking conclusion of this work is that eq. (2) is constructed entirely within the approximation of *infinite* population size, yet it recovers fitnesses from noisy, *finite*-population synthetic and experimental datasets accurately. For decades, the genetic drift term in eq. (1) has been a hallmark of classical population genetics, and it generally cannot be ignored in any forward-time predictive modeling of evolution. However, we have found in this work that its exclusion allows for simple mathematical manipulation of eq. (1) into eq. (2), permitting both ease of use as well as high accuracy fitness inference from time series without any restrictive approximations on the order of epistasis.

Looking forward, we anticipate that eq. (2) will play an important role in computational *fitness landscape design (FLD)* (Mohanty and Shakhnovich 2025, 2026), which includes recently introduced methods for quantitatively tuning evolutionary fitness landscapes, with downstream applications optimal suppression of fitness-increasing viral mutations using designed antibodies. We suggest that eq. (2) will be useful in “closed-loop” FLD: antibody sequences can be generated, creating new effective fitness landscapes, viral evolution can take place in the presence of the generated antibody, and FLD-imposed fitnesses can be estimated using eq. (2) in one large computational loop to search for optimal antibodies which best suppress viral fitness growth.

## Materials and Methods

### Numerical simulations

#### Random epistatic fitness landscapes

For the Wright-Fisher, Moran, and ProSeD simulations, we generated random fitness landscapes with epistasis. We used a binary genomic alphabet with size 2 and sequence length 10, so there were 1024 genotypes in total. For Wright-Fisher and Moran simulations, epistatically sparse fitness landscapes were generated—to emulate realistic epistatic sparsity of empirical fitness landscapes (Poelwijk *et al*. 2019; Aghazadeh *et al*. 2021; Brookes *et al*. 2022; Tsui *et al*. 2026). First, we chose a mean sparsity fraction of *ρ* = 0.05. Then we constructed a random vector (corresponding all 1024 possible epistatic coefficients of the fitness landscape) of length 1024 where each position was masked with probability 1 − *ρ* = 0.95. This vector was multiplied a random vector where each element was drawn i.i.d from a normal distribution *N* (0, 0.01). We call this product of the mask vector and the random vector **J**, the sprase vector of all (signed) epistatic coefficients for the fitness landscape. We use the (inverse) Walsh-Hadamard transformation to obtain the fitness landscape **X** = *H***J**, where *H* is proportional to a Hadamard matrix.

For ProSeD simulations, we assigned each of the 1024 genotypes *g* a replication probability *π*_*g*_ (per microscopic time step) i.i.d. from a beta distribution *π*_*g*_ ∼ Beta(2, 5). We set the number of offspring per replication event to *m* = 2, and we set the number of microscopic time steps between broth dilutions to Δ*t*_*b*_ = 4. The fitness of genotype *g* is then approximately *X*_*g*_ ≈ *π*_*g*_*m*/*τ*.

#### Wright-Fisher simulations

We follow the version of the Moran process outlined by Greenbury *et al*. (2022). A population of size *N* was initialized at a random starting genotype. For the selection step, the next generation was selected by sampling, for each individual with replacement, from the previous generation’s individual with probability weights proportional to 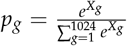

For the mutation step, each genomic site for each newly selected individual’s could then be mutated with probability *µ* = 0.01. The process was repeated for *T* user-specified generations.

#### Moran simulations

We follow the version of the Moran process outlined by Good (2024). A population of size *N* was initialized at a random starting genotype. For each microscopic time step, two individuals, *a* and *b*, were randomly chosen from the population. Individual *a* was selected as the winner from a Bernoulli trial with probability 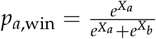. If *a* wins, then *a* returns to the population and also replaces *b*. If *b* wins, then *b* returns to the population and also replaces *a*. This procedure was repeated a total of *TN* times for *T* user-specified macroscopic time steps. Frequencies were recorded every *N* time steps (corresponding to *N* pair challenges).

#### ProSeD simulations

Following Mohanty *et al*. (2026), a population of size *N* was initialized at a random starting genotype. For each microscopic time step, individuals were chosen to replicate as i.i.d. Bernoulli trials with probabilities given by the replication probabilities *π*_*g*_ obtained from the beta distribution previously. Individuals chosen to replicate produced *m* offspring with the same genotype, and the original individuals were deleted from the population. Each genomic site for each individual could then be mutated with probability *µ* = 0.01. This process was repeated a total of *τ* = 4 times, after which the population was down-sampled (without replacement) to *N*, and genotype frequencies were recorded after downsampling.

#### Fit-Seq2.0 Growth Simulations

The code from Li *et al*. (2023) was directly run from the command line with the following parameters: time between broth transfers: 4, average read depth: 100, genomic DNA copies: 500 (default), PCR cycles: 25 (default).

## Code and data availability

Code and simulated data will be made public upon publication. For barcoded yeast experiments, time series and author-estimated fitnesses are publicly available from Nguyen Ba *et al*. (2019). SARS-CoV-2 pathogen genomic frequency data and fitnesses from Obermeyer *et al*. (2022) were collated in Wang *et al*. (2024). For MNV-1 evolution experiments, time series are publicly available from Rotem *et al*. (2018), and fitness estimation can be performed with the code posted by Mohanty and Shakhnovich (2026).

## Acknowledgments and Funding

This work was supported by a Hertz Foundation Fellowship (to V.M.), a Paul and Daisy Soros Fellowship (to V.M.), and by award T32GM14427 from the National Institute of General Medical Sciences (to Harvard/MIT MD-PhD Program). The content is solely the responsibility of the authors and does not necessarily represent the official views of the National Institute of General Medical Sciences or the National Institutes of Health. We thank the MIT Engaging Cluster, supported by the MIT Office of Research Computing and Data, for computational resources.

## Supplementary Information

### S1 Fitness can be estimated from genotype correlation time series data

Consider a population of *N* haploid, asexually reproducing organisms. The forward-time, continuous-time evolution of frequency *f*_*g*_ of a genotype *g* is given by the stochastic differential equation:

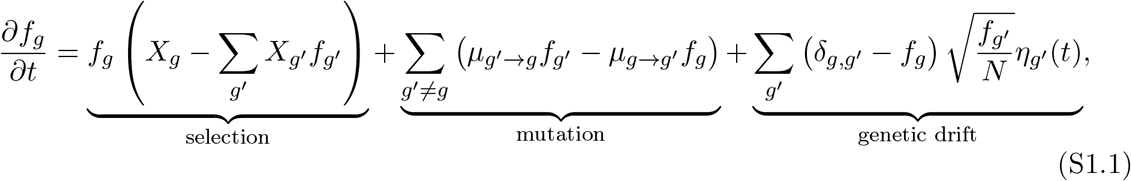

where *X*_*g*_ is the fitness of genotype *g, µ*_*g*→*g*_′ is the mutation rate from genotype *g* to *g*^′^, and *η*_*g*_′ (*t*) is a Gaussian noise term with *η*_*g*_′ (*t*) = 0 and *η*_*g*_(*t*)*η*_*g*_′ (*t*^′^) = *δ*_*g,g*_′ *δ*(*t* − *t*^′^). In the limit of large population (*N* → ∞), we can approximate the diffusion term to be small relative to the selection and mutation terms. These operations cause the above equation to now become

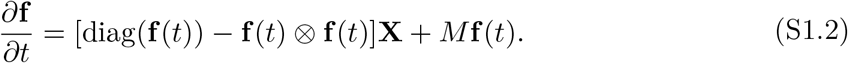

We now perform time averaging to obtain

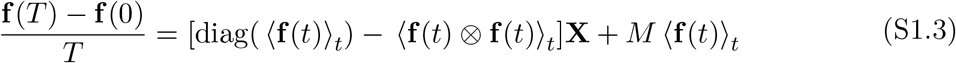

where 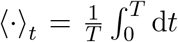 denotes time averaging, and we have defined a mutation rate matrix *M*_*gg*_′ = *µ*_*g*_′_→*g*_ for *g*≠ *g*^′^ and *M*_*gg*_ = − ∑_*g*_′ _*g*_ *µ*_*g*→*g*_′ . Recognizing that absolute mutation rates may be affected by time discretization, we use a rescaled relative mutation rate matrix 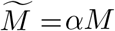, with *α* being a scaling factor. Similarly, only relative fitnesses affect the evolutionary dynamics, so the fitnesses **X** should absorb any scaling factor. Letting 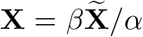, choosing *β/α* as a scaling factor, and rearranging into a matrix equation, we have

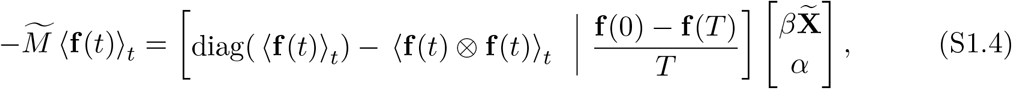

The estimated relative fitness landscape 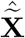 can be obtained from linear regression by minimizing the least squares loss of the right and left sides of the equation, above, computable with the pseudoinverse

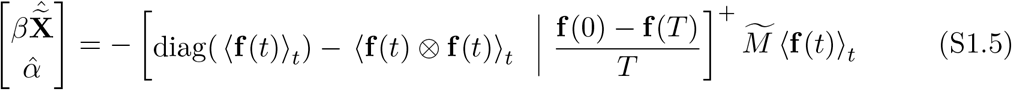

The Moore-Penrose pseudoinverse form *A*^+^ = (*A*^*T*^ *A*)^−1^*A*^*T*^ does not exist because (*A*^*T*^ *A*)^−1^ is singular, since *A* has one more column than rows. But, the pseudoinverse always exists and can be calculated with singular value decomposition (SVD). If *A* = *U* Σ*V* ^*T*^ is the SVD of *A*, then *A*^+^ = *V* Σ^−1^*U*^*T*^ , which is approximated by 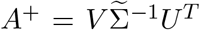, where 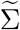 has the smallest singular values removed.

### S2 Alternate equation for exactly known mutation rate matrix

In the infinite population limit *N* → ∞, we have

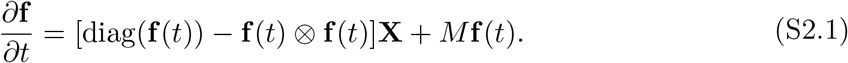

We now perform time averaging to obtain

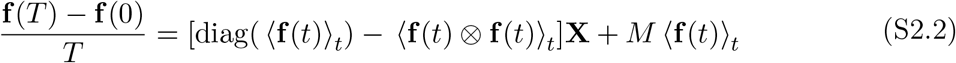

where 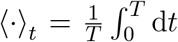 denotes time averaging, and we have defined a mutation rate matrix *M*_*gg*_′ = *µ*_*g*_′_→*g*_ for *g* ≠ *g*^′^ and *M*_*gg*_ = − ∑_*g*′≠*g*_ *µ*_*g*→*g*_′ . When the mutation rate is exactly known (with respect to the correct time coarse-graining), a modified inference protocol can be used which does not involve rectangular matrices. We can rearrange the above equation to obtain We now perform time averaging to obtain

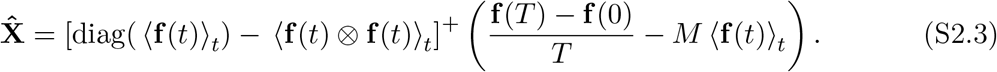

The pseudoinverse becomes an inverse when the matrix is invertible.

### S3 Alternate equation in the strong selection, weak mutation limit

In the strong selection, weak mutation (SSWM) limit, mutation rates are typically much slower than fitnesses (growth rates). As a result, the evolutionary dynamics are typically dominated by selective sweeps in which one genotype fixes in the population before the next mutation is introduced. This leads to principally monomorphic dynamics in which the entire population tends to be concentrated at one genotype, then another genotype, then another, and so on.

Like in the main text, we recognize that absolute mutation rates may be affected by time discretization, we use a rescaled relative mutation rate matrix 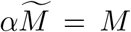, with *α* being a scaling factor (note that the convention is slightly different from the rescaling in the main text). We again rearrange eq. (S2.2) into a matrix equation, but including the mutation term in the matrix to be pseudoinverted. This leads to an equation that is the same as what we presented in the main text, but with the boundary term and the mutation term swapped:

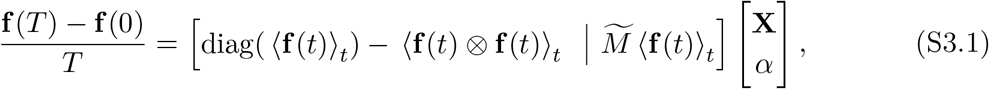

The estimated relative fitness landscape 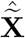 can be obtained from linear regression by minimizing the least squares loss of the right and left sides of the equation, above, computable with the pseudoinverse

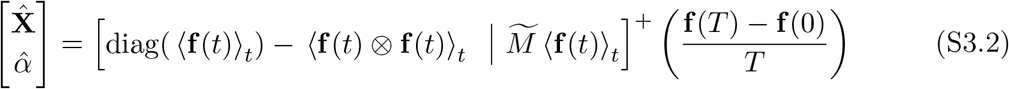

Once again, the exact pseudoinverse *A*^+^ = (*A*^*T*^ *A*)^−1^*A*^*T*^ does not exist because (*A*^*T*^ *A*)^−1^ is singular. If *A* = *U* Σ*V* ^*T*^ is the SVD of *A*, then *A*^+^ = *V* Σ^−1^*U*^*T*^ , which is approximated by 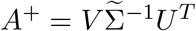, where 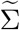 has the smallest singular values removed.

#### S3.1 Zero Mutation Matrix

When truly zero mutations are present, the mutation matrix vanishes, so only the derivative term 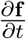 can contribute to the inference process. In this case, we simply use eq. (S3.3), but since *M* = 0, we can simply remove the extra column from the pseudoinverted matrix (and the *α* parameter is thus arbitrary):

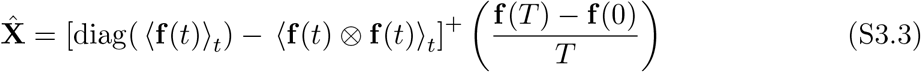

#### S3.2 Proof of exactness of Equation (S3.3) for a sequence of selective sweeps

We now analytically consider the case of *n* selective sweeps occurring in an infinite population hopping from genotype to genotype in a sequence of increasing fitness. We will prove that eq. (S3.3) yields fitness estimates that have Pearson correlation *r* = 1 with the original fitnesess.

Consider a set of genotypes labeled by indices {1, … , *n*} and their fitnesses *X*_1_ *< X*_2_ *<* · · · *< X*_*n*_. In our scenario, *f*_1_(0) = 1 − *ϵ* (for 0 *< ϵ <* 1*/*2) fraction of the population begins at genotype 1 at time *t* = 0 while the remaining *ϵ* fraction of the population is on genotype 2, so *f*_2_(0) = *ϵ*. As a result, over a time window *T*_1→2_, genotype 2 will fix in the population and the fraction of population 1 will become small. In particular, we will define *T*_1→2_ to be so that at some point infinitesimally before time *t* = *T*_1→2_, fraction of genotype 1 becomes *f*_1_(*T*_1→2_) = *ϵ* and 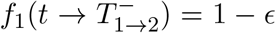. At this time, a very rare mutation occurs in which genotype 1 disappears (this can also follow from assuming that *N* is large but finite) and genotype 3 appears at frequency 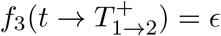. Another selective sweep then occurs over a time *T*_2→3_ in which genotype 3 fixes in the population, sending the frequency of genotype 2 eventually to *ϵ*. This procedure repeats in a sequence of increasing fitness so that ultimately the time series ends with *f*_*n*_(*T*) = 1 − *ϵ* and *f*_*n*−1_(*T*) = *ϵ*, with 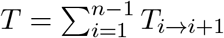. At all times *t*, only two genotypes have nonzero frequencies.

In the infinite population limit, we can write the diffusion limit of population genetics as a replicator equation

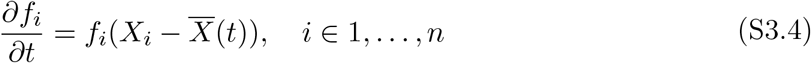

with 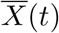 representing the mean fitness of the population. If we a consider a time segment *T*_*i*→*i*+1_ for *i* ∈ {1, … , *n* − 1}, the mean population only involves two frequencies:

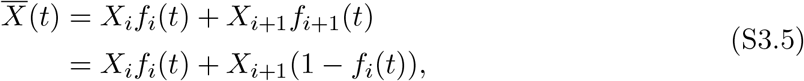

so we have

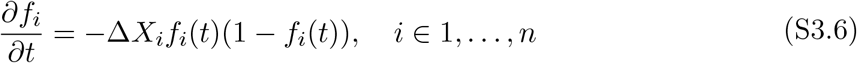

where Δ*X*_*i*_ = *X*_*i*+1_ − *X*_*i*_ *>* 0 and is defined for *i* ∈ 1, … , *n*. Equation (S3.6) has an exact solution:

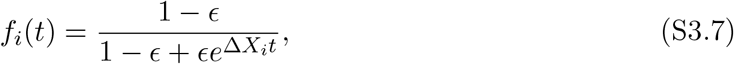

where we have, without loss of generality, shifted the time so that the start of the sweep is at *t* = 0 (even if *i >* 1). By setting *f*_*i*_(*T*_*i*→*i*+1_) = *ϵ*, we can find the time interval *T*_*i*→*i*+1_ over which the sweep occurs.

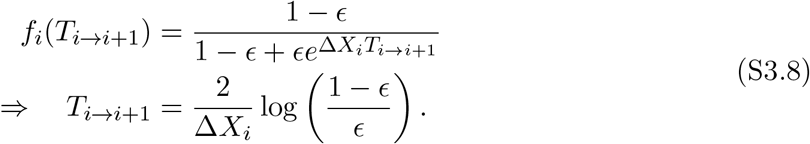

In the interval *T*_*i*→*i*+1_, the the integral of the frequency is its average (1/2, by symmetry) times the time duration

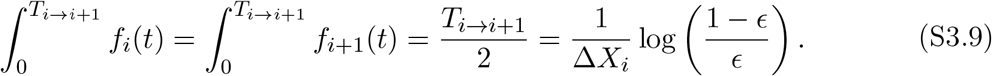

The integral of the frequency squared is exactly computable (both genotypes *i* and *i* + 1 have the same value of the integrated frequency squared, by symmetry):

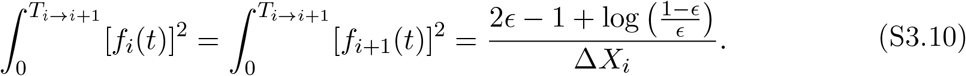

We now must compute the elements the matrix in eq. (S3.3). We first start with the general *i* ∈ 2, … , *n* − 1, excluding the first and last genotypes. Each average will be built from contributions from two time segments, *T*_*i*−1→*i*_ and *T*_*i*→*i*+1_, which will be the sum of two of the integrals computed above:

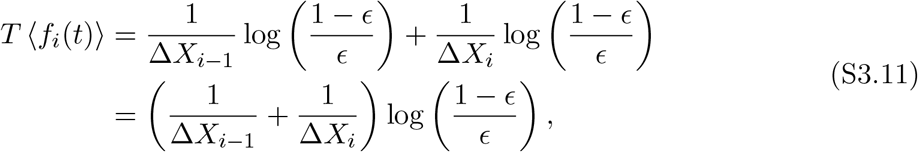

and

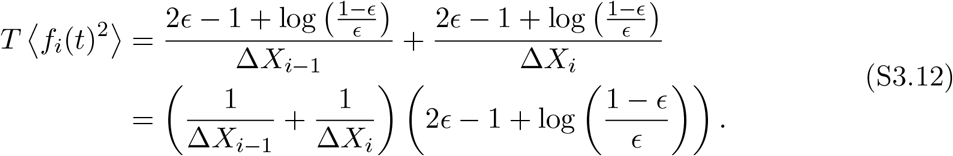

Thus, we can get the diagonal elements of the matrix for *i* ∈ {2, … , *n* − 1}:

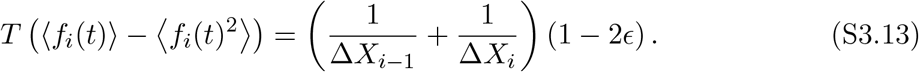

For *i* ∈ {1, … , *n* − 1}, the off-diagonal elements are also simply computable:

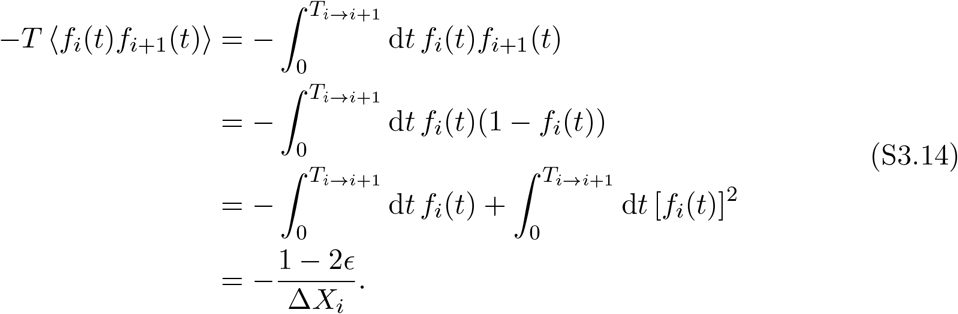

The last two terms left to calculate are the diagonal elements for genotypes 1 and *n*:

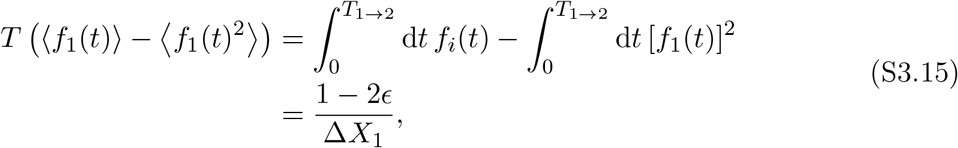

and

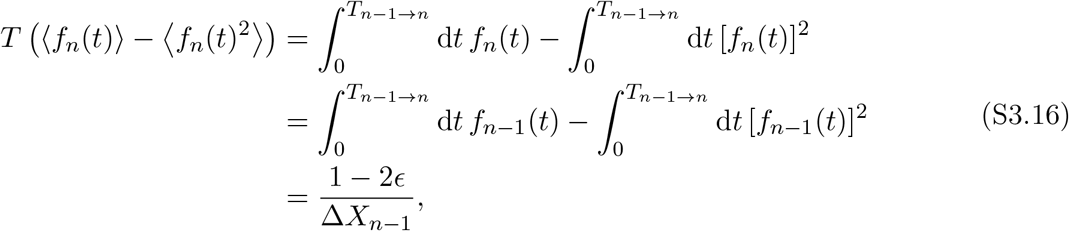

where in the second equality we have used the symmetry of the trajectories.

We now define the matrix

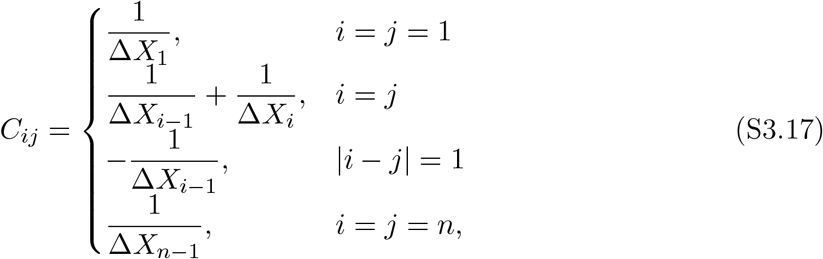

which is a tridiagonal, symmetric matrix. Bringing out a factor of 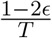, we have

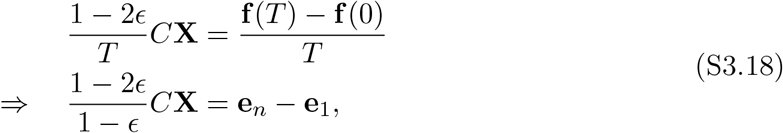

where **e**_*i*_ is a unit vector pointing in the *i*-th direction. Absorbing the constant 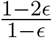 into the fitness vector 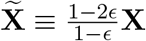, we have

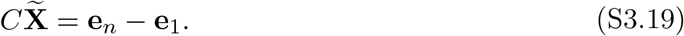

Computing the exact pseudoinverse *C*^+^ is difficult, but we can recall that *C*^+^(**e**_1_ − **e**_*n*_) will be provide the minimum least squares solution 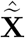 with the smallest *ℓ*_2_ norm, minimizing

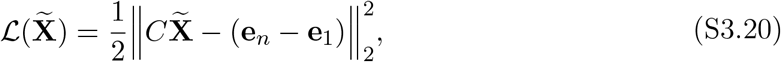

which may not have a unique solution. Using these two facts of the pseudoinverse, we will now exactly compute *C*^+^(**e**_1_ − **e**_*n*_).

First, we show that the original fitness vector **X** minimizes the least squares loss. First, we compute *C***X**:

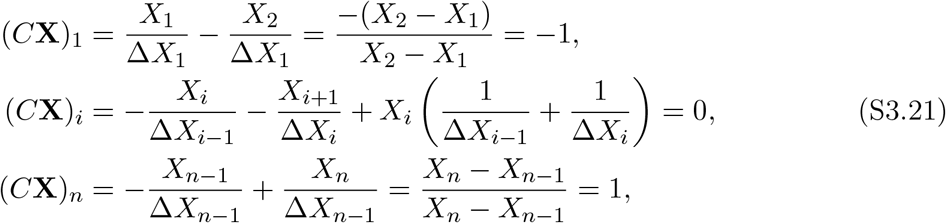

where *i* ∈ {2, … , *n* − 1}, so we can write

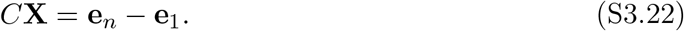

Thus, 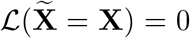. Since 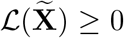, we know that 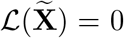 minimizes the least squares loss. However, we do not know if this solution is unique, and if it not unique, then which solution would be returned by 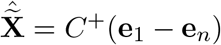.

Now, we prove that the solution is not unique because *C* is not invertible. We show this by noting that 1, the vector of all ones, is in the kernel of *C*:

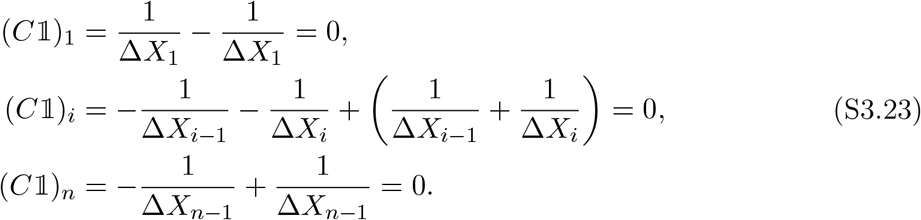

so we can write

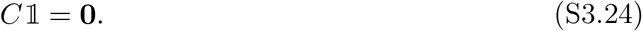

Thus, *C* is rank-deficient and therefore singular. Additional least squares-minimizing solutions can thus be constructed by adding any multiple of the homogeneous solution 1 and the particular solution **X**. This makes physical sense because the original replicator equations are invariant to constant shifts in the fitness.

We now must show that the space of all possible solutions has been found and that there are no other homogeneous solution. We do this by showing that rank(*C*) = *n* − 1. Noting that *C* is a symmetric tridiagonal matrix, we first define an (*n* − 1) × (*n* − 1) matrix *B* such that

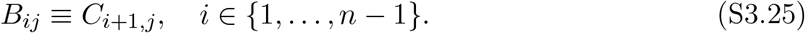

*B* is an upper triangular matrix whose diagonals are known. The determinant of an upper triangular matrix are given by the product of its diagonal elements, so we have

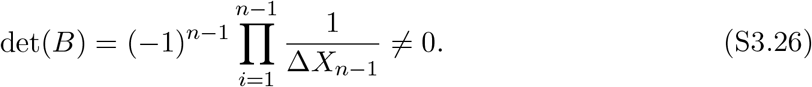

Since the determinant of *B* is nonzero, we know that *B* is full rank. Now, since *B* is a submatrix of *C*, we must have that rank(*C*) ≥ rank(*B*) = *n* − 1. Since we have found a homogeneous solution to 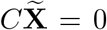, it immediately follows that rank(*C*) ≤ *n* − 1. Together, this means we have rank(*C*) = *n* − 1.

Therefore the set of all least squares-minimizing solutions is given by a linear combination of **X** + *λ*1. We now must find 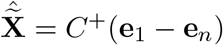, which will be given by the solution **X** + *λ*1 with *λ* chosen such that 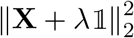 is minimized. We do this by direct optimization:

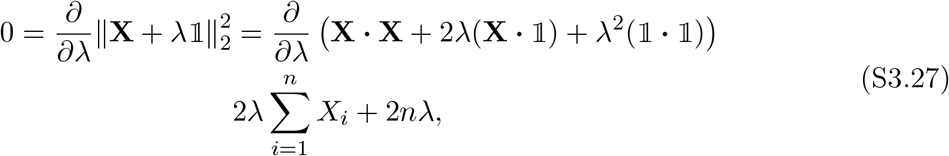

from which it follows that

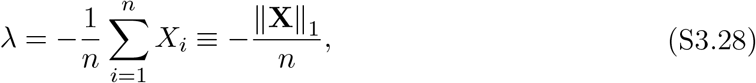

which is the negative of the mean of the entire fitness vector.

We now have that the exact inferred fitness vector given by eq. (S3.3)

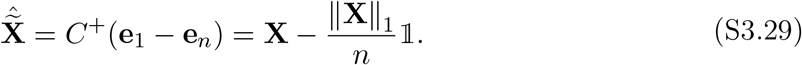

Indeed, the Pearson correlation *r* = 1 since the inferred fitness vector 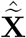 and true fitness vector **X** are linearly related. The proof is complete.

### S4 Derivation of negative identity matrix approximation for scaled mutation matrix for large genotype spaces

Suppose there are *G*_poss_ genotypes that can be observed, and *G*_obs_ genotypes that are actually observed. Suppose we define the mutation matrix *M* to be a *G*_poss_ × *G*_poss_ matrix on the genotype space of all *possible* genotypes, not necessarily just the observed ones; accordingly, ⟨**f** (*t*)⟩_*t*_ is a vector of length *G*_poss_ corresponding to the time-averaged frequency of all possible genotypes, observed and unobserved.

The mutation rate matrix has the same definition as before: *M*_*gg*_′ = *µ*_*g*_′_→*g*_ for *g*≠ *g*^′^ and *M*_*gg*_ = − ∑_*g*_′ _*g*_ *µ*_*g*→*g*_′ . We had also defined a rescaled relative mutation rate matrix 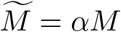, with *α* being a scaling factor that does not have to be known *a priori*. In Equation (S3.3), the matrix 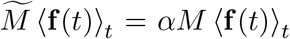 appears in the computation. For a genotype *g*, we can write an element of the vector

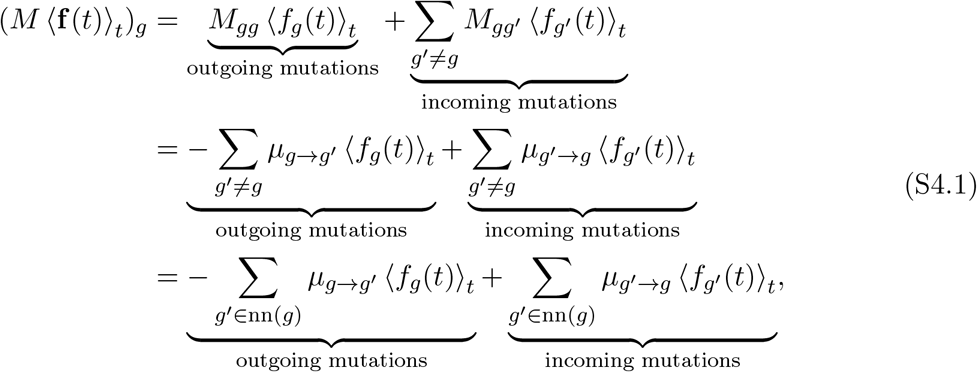

where we have used nn(*g*) to denote the *set* of genotypes which are nearest neighbors of genotype *g*. Also, let *n*_poss_(*g*) ≡ |nn(*g*)| be the *number* of possible outgoing mutational neighbors from genotype *g*.

Now, suppose that *µ*_*g*→*g*_′ and *µ*_*g*_′_→*g*_ are only nonzero if genotypes *g* and *g*^′^ are neighbors in the genotype space (e.g. accessible via a point mutation). We then work in the approximation that a genotype mutation from *g* to any neighboring genotype *g*^′^ happens with probability *µ*_*g*→*g*_′ ≡ *µ/n*_poss_(*g*). This means that any outgoing mutation has uniform probability of going from genotype *g* to any of its *n*_poss_(*g*) neighbors with uniform probability, and the total outgoing mutational probability from any genotype *g* is

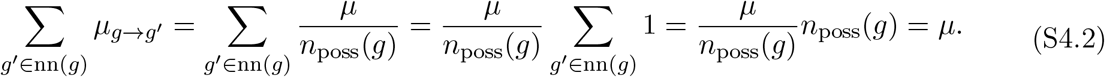

Therefore, we can also write

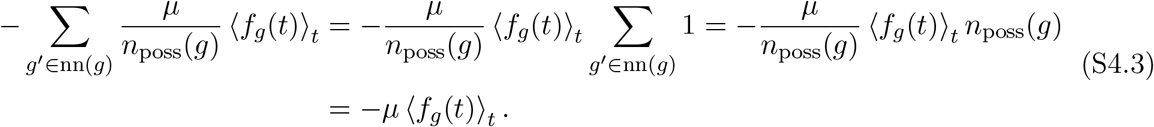

For incoming mutations, we can write

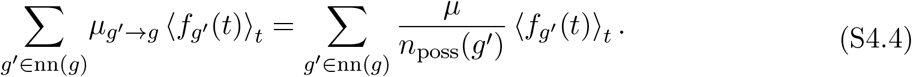

If we assume that the number of neighbors is the same for every genotype (as is the case for fixed genome lengths with fixed alphabets), then we can further write

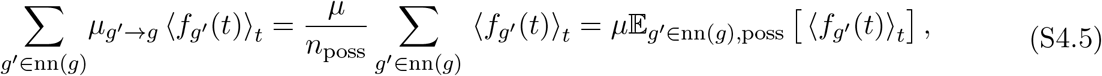

where E_*g*_′_∈nn(*g*),poss_ ⟨*f*_*g*_′ (*t*)⟩_*t*_ is average of the mean frequencies of all neighbors of genotype *g*. Then, the diagonal term in the mutation matrix dominates over the off-diagonals, for a genotype *g*, if

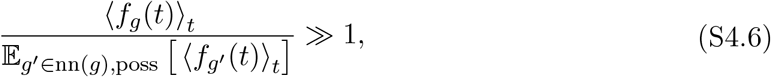

which means the observe frequency of genotype *g* should greatly exceed the frequencies of its neighbors. We argue that in very large genotype spaces, the neighbor-averaged mean frequency E_*g*_′_∈nn(*g*),poss [_⟨*f*_*g*_′ (*t*)⟩_*t*_] will strongly be skewed by a large number of zeros due to the vast majority of genotypes going unobserved; thus, the condition eq. (S4.6) should be valid for experimental settings like barcode experiments and microbial evolution experiments with long-read sequencing.

To strengthen the argument and provide better intepretation for eq. (S4.6), we can continue: writing *n*_obs_(*g*) as the number of *observed* neighbors of genotype *g*, we can also rewrite the expectation E_*g*_′_∈nn(*g*),poss [_⟨*f*_*g*_′ (*t*)⟩_*t*_]

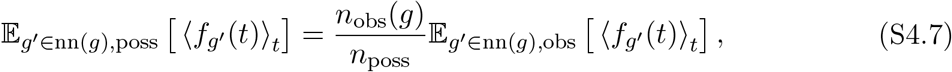

where E_*g*_′_∈nn(*g*),obs [_⟨*f*_*g*_′ (*t*)⟩]_*t*_ is the expectation of average frequency of the observed neighboring genotypes of genotype *g*. This yields an equivalent condition

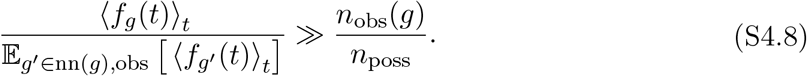

Again, the ratio of observed neighbors *n*_obs_(*g*)*/n*_poss_ is likely to be very small *n*_obs_(*g*)*/n*_poss_ ≪ 1, and is guaranteed to be less than or equal to 1: *n*_obs_(*g*)*/n*_poss_ ≤ 1. Then, if genotype *g* is a fitness peak, then it is more likely to be selected over time and more abundant in the time series compared to its neighbors, so we would expect 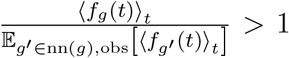. This, in addition to sparse observation of the vast genotype space (*n*_obs_(*g*)*/n*_poss_ ≪ 1), would suggest that eq. (S4.8) should hold. If genotype *g* has its frequency close to the average of its neighbors (for example, possibly because *g* is a member of a neutral set or otherwise has comparable fitness to its neighbors), then 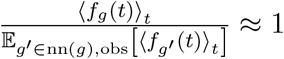. and eq. (S4.8) still holds because of the sparsity of the observed subset of the genotype space. Only if genotype *g* is rare compared to all of its observed neighbors would we have 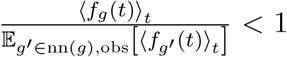. This could occur, for instance, if *g* occurs in a sharp fitness valley. Even then, an additional condition is required; namely, that many of genotype *g*’s possible neighbors are observed: 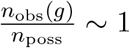. Only then is eq. (S4.8) likely to be violated.

Based on this interpretation, we believe that for large genotype spaces, eq. (S4.6) (and, equivalently, eq. (S4.8)) is likely to be true, which would mean that contributions to each element of *M* ⟨**f** (*t*)⟩_*t*_ from outgoing mutations will outweigh the sum of contributions from incoming mutations:

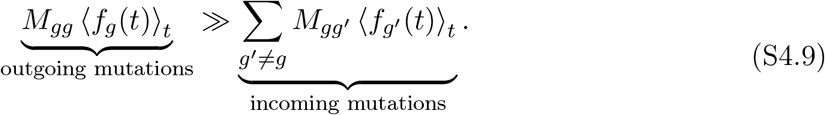

Then, it follows that

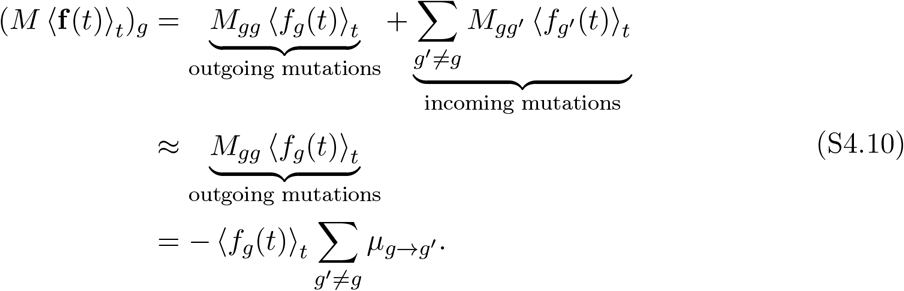

Using *µ*_*g*→*g*_′ = *µ/n*_poss_(*g*), this simplifies to

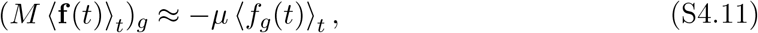

or

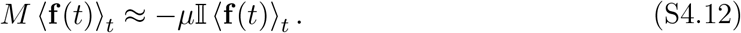

When using the scaled mutation matrix 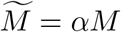, we can simply absorb the mutation rate *µ* into the positive constant *α*, writing

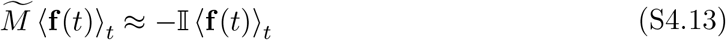

In the fitness estimation equation eq. (S1.5), the mutation matrix only appears in the context of acting as an operator on the average frequency vector. Therefore, we can replace 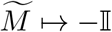 when the mutation matrix is not known or difficult to calculate, and if the genotype space is very large. In the Main Text we show that this works well for barcoded cell growth simulations, two real barcoded yeast evolution experiments, and two murine norovirus 1 serial evolution experiments which use long-read sequencing.

## S5 Supplementary Figures

**Figure S1.**
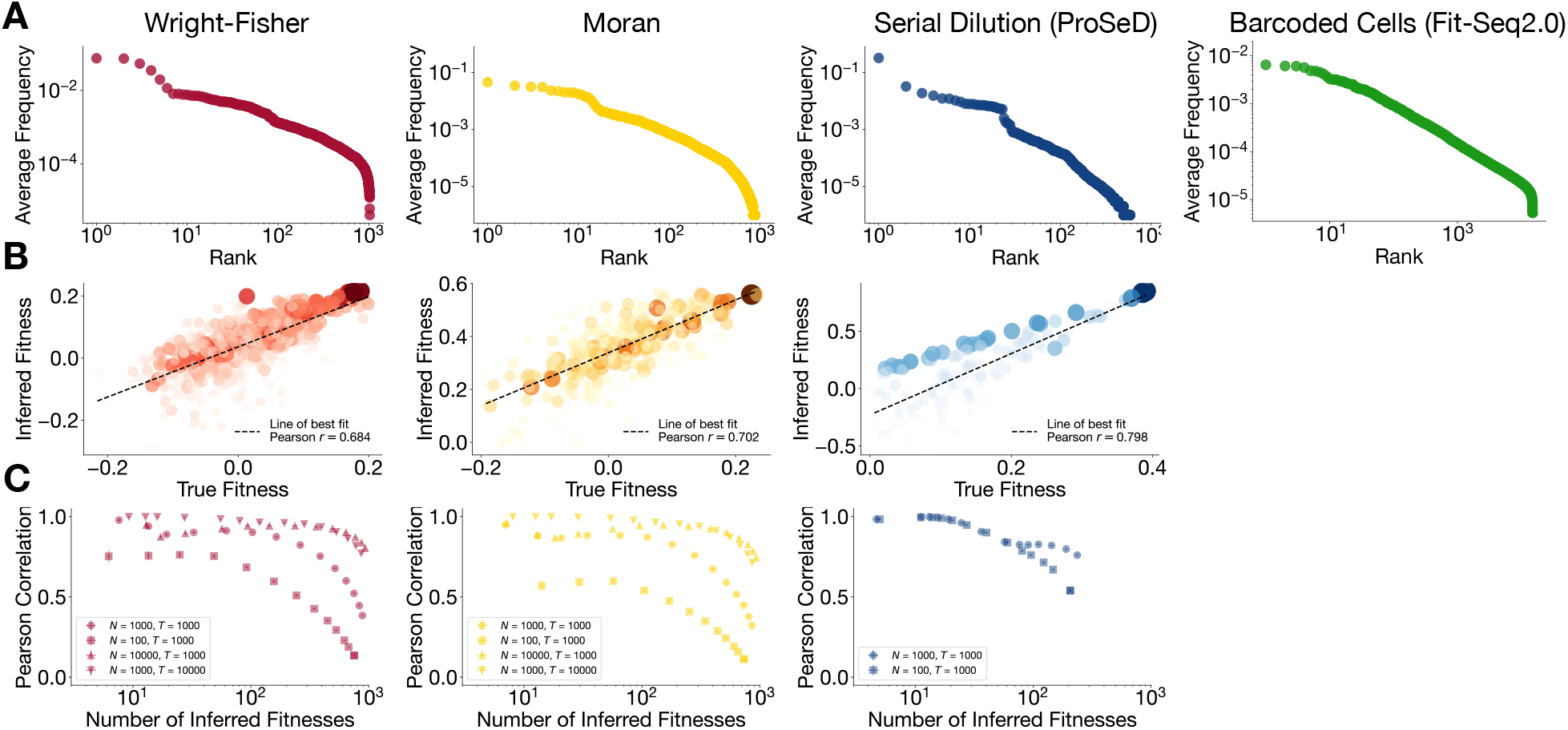
Extended data for Main Text Figure 2: Fitness inference for simulated evolutionary dynamics using Main Text eq. (2). (**A**) Average genotype frequency versus rank for Wright-Fisher, Moran, ProSeD, and Fit-Seq2.0 Growth Simulations, demonstrating ≳ 4 orders of magnitude variation in genotype average frequency. (**B**) Comparison between ground truth fitnesses and fitnesses inferred using Main Text eq. (2) on example trajectories for each simulation type, evaluating on genotypes whose average frequency was ⟨*f* (*t*)⟩_*t*_ ≥ 10^−4^ . Main Text eq. (2) demonstrates faithful reconstruction of the observed fitnesses even for rare genotypes. (**C**) Pearson correlation between ground-truth fitnesses and fitnesses inferred using Main Text eq. (2), as a function of number of inferred fitnesses, which is set by the average frequency cutoff. Multiple population sizes and/or simulation lengths are shown. Error bars are standard errors.

**Figure S2.**
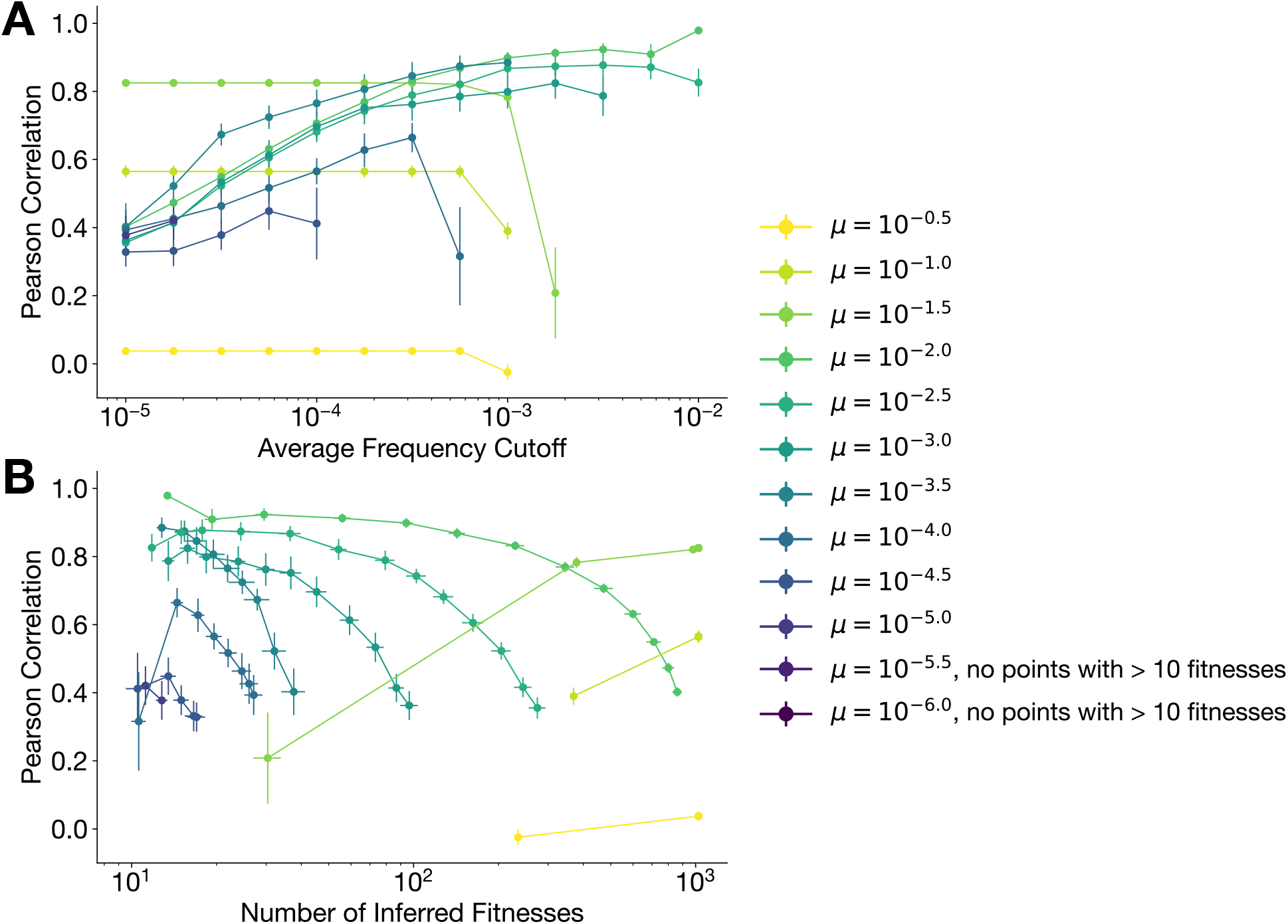
Pearson correlation of recovered and true fitness landscapes for varied mutation probabilities (per-site, per-individual, per generation) in Wright-Fisher simulations. (**A**) Pearson correlation between fitnesses inferred using Main Text eq. (2) and literature-reported fitnesses, as a function of number of inferred fitnesses, as a function of average frequency cutoff. (**B**) Pearson correlation between fitnesses inferred using Main Text eq. (2) and literature-reported fitnesses, as a function of number of inferred fitnesses, as a function of number of inferred fitnesses, which is set by the average frequency cutoff. Curves are plotted for simulations with different mutation probabilities; points are averaged over 10 trials. All curves are plotted for simulations with different mutation probabilities; points are averaged over 10 trials; error bars are standard errors.

**Figure S3.**
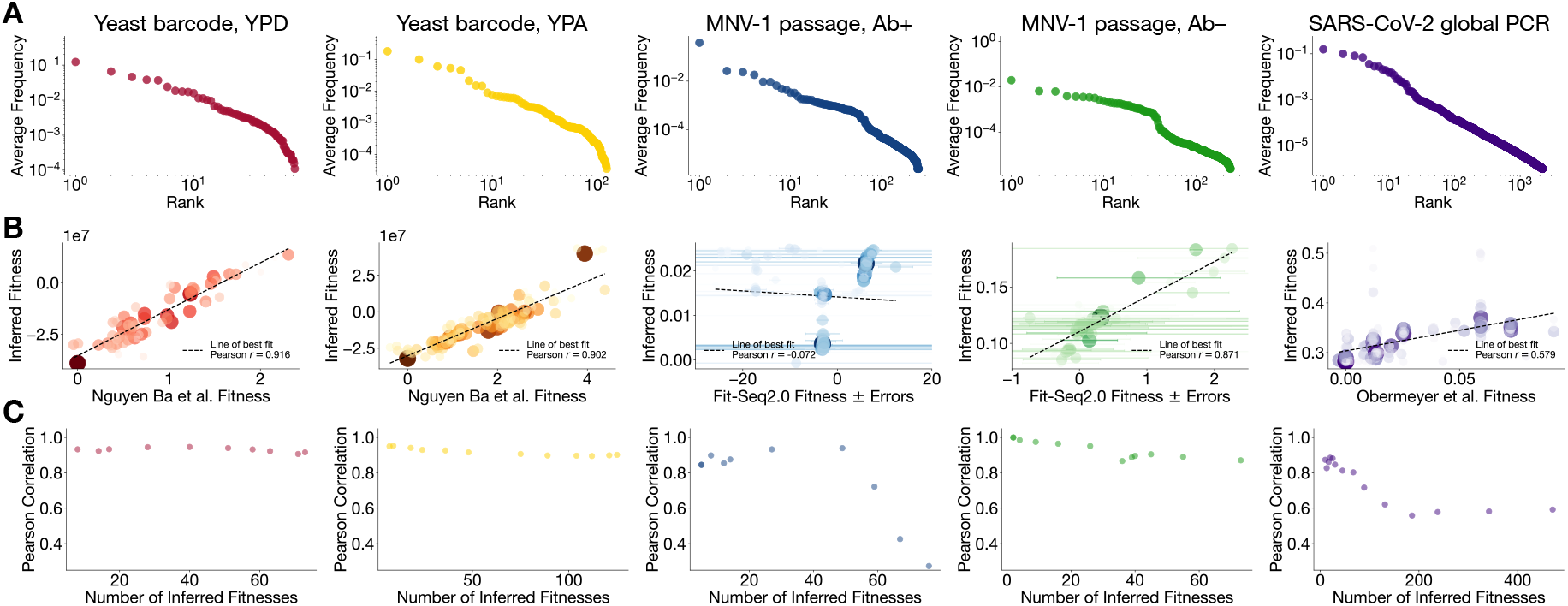
Extended data for Main Text Figure 3: Fitness inference for empirical evolutionary dynamics using Main Text eq. (2). (**A**) Average genotype frequency versus rank each empirical time series, demonstrating ≳ 4 orders of magnitude variation in genotype average frequency. (**B**) Comparison between fitnesses inferred using Main Text eq. (2) and fitnesses inferred by other methods in the literature; we included genotypes whose average frequency was ⟨*f* (*t*)⟩_*t*_ ≥ 10^− 4.5^ . Main Text eq. (2) demonstrates reconstruction of the observed fitnesses comparable to those found by other methods. (**C**) Pearson correlation between fitnesses inferred using Main Text eq. (2) and literature-reported fitnesses, as a function of number of inferred fitnesses, which is set by the average frequency cutoff.

